# Functional redundancy in natural pico-phytoplankton communities depends on temperature and biogeography

**DOI:** 10.1101/2020.05.14.096123

**Authors:** Duyi Zhong, Luisa Listmann, Maria-Elisabetta Santelia, C-Elisa Schaum

## Abstract

Biodiversity affects ecosystem function, but how this relationship will pan out in a changing world is still a major question in ecology. It remains especially understudied for pico-phytoplankton communities, which contribute to carbon cycles and aquatic food webs year-round. Observational studies show a link between phytoplankton community diversity and ecosystem stability, but there is only scarce causal or empirical evidence. Here, we sampled phytoplankton communities from two biogeographically distinct (but close enough to not be confounded by differences in day length and precipitation) regions in the Southern Baltic Sea, and carried out a series of dilution/regrowth experiments across three assay temperatures. This allowed us to investigate the effects of loss of rare taxa and establish causal links in natural communities between species richness and several ecologically relevant traits (e.g. size, biomass production, and oxygen production), depending on sampling location and assay temperature. We found that the samples’ bio-geographical origin determined whether and how functional redundancy changed as a function of temperature for all traits under investigation. Samples obtained from the slightly warmer and more thermally variable regions showed overall high functional redundancy. Samples from the slightly cooler, less variable, stations showed little functional redundancy, i.e. function decreased the more species were lost from the community. The differences between regions were more pronounced at elevated assay temperatures. Our results imply that the importance of rare species and the amount of species required to maintain ecosystem function even under short-term warming (e.g. during heat waves) may differ drastically even within geographically closely related regions of the same ecosystem.

## Introduction

When ecosystems lose species, the function and services of the ecosystem may decline. The insurance hypothesis states that ecosystem stability increases with biodiversity [1]. Evidence has long been emerging for this to be true across taxa and biomes, spanning heterotrophic and autotrophic microbial communities to plants to metazoans, i.e. ecosystems with higher biodiversity tend to be more productive on average regardless of the type of organism under investigation [2-6]. In a warming world, variation in temperature may shape the biodiversity-ecosystem function relationship on short and long-term time scales [7,8]. A recent study has experimentally examined the synergistic effects of warming and biodiversity loss on function in bacterial assemblages [9], but no comparable data exist for natural phytoplankton communities. As a consequence, studies on phytoplankton biodiversity and ecosystem function remain largely observational [2] and usually do not consider loss of biodiversity in interaction with aspects of climate change (but see [10]) or evolutionary history (but see [11] for population genetics). Pico-phytoplankton contribute about 20% of phytoplankton primary production [12], and unlike larger phytoplankton, they do not form blooms, but contribute to the photosynthesising foundation of aquatic ecosystems all year [13,14]. They show rapid physiological [15,16] and evolutionary [17,18] responses to changing environments when studied as single-strain cultures. There is isolated evidence that the speed and mechanism of evolutionary responses in a changing environment may differ between phytoplankton strains evolving in isolation and strains evolving in communities, mixed cultures, or multi-genotype biofilms [19-23]. Comparable experiments have not been carried out for pico-phytoplankton (but see [24]). As a consequence, we need to conduct manipulative experiments to establish a better understanding of how the links between pico-phytoplankton community function and pico-phytoplankton community diversity change with temperature on short- and long-term timescales. Since natural phytoplankton strains are notoriously difficult to grow in isolation under laboratory conditions, assembling artificial communities from natural phytoplankton components can be an arduous task. Organisms from culture collections which have already been shown to grow well in isolation are easier to assemble into communities (e.g.[10]) but may not reflect well the complex environments that the lineages were originally sampled from. Therefore, when working with natural assemblages, serial dilution of natural samples can provide a useful tool to reduce biodiversity to such an effect that rare species are lost, and common species remain [25,26].

Here, we investigated the combined effects of transient warming and biogeographic history on the diversity-function relationship in natural pico-phytoplankton communities. To do so, we obtained pico-phytoplankton community samples during two cruises of RV ALKOR (AL505 and AL507) on the Southern Baltic Sea in 2018 (see Figure 1 and Table S1). Of the two basins sampled, the Kiel Bight is characterised by, on average 2 °C higher temperature than the Bornholm Basin and unpredictable fluctuations in temperature on the timescale of days to months (Santelia *et al*. in prep – SI document 2). In the Bornholm Basin, fluctuations in temperature follow a highly predictable pattern governed by seasonality (Santelia *et al*. in prep – SI document 2). The basins are connected through currents [27], and the regions are geographically close to each other (see Figure 1) so that we can rule out confounding effects introduced by e.g. differences in precipitation, day length, and light intensity [28]. By assaying the effect of species loss on communities across three temperatures within the range of Southern Baltic Sea spring and summer temperatures [27,28], we can test the contributions of long-term (comparison of basins) and short-term (assay temperatures) changes in temperature on key community functions.

**Figure 1:**
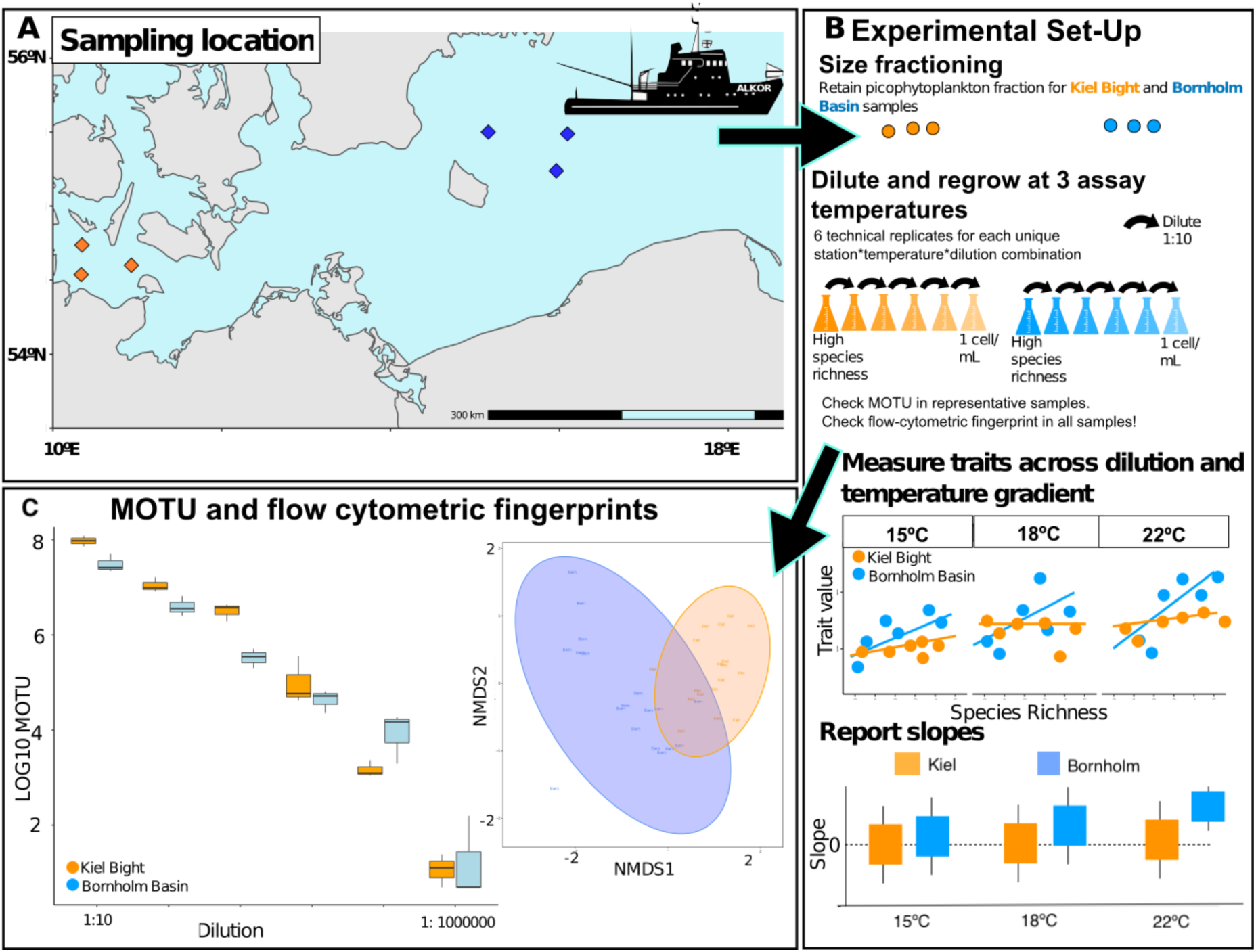
Overview of sampling locations, experimental set-up, and assessment of species richness via meta-barcoding and flow cytometry. A) sampling locations: We took samples in two biogeographically distinct regions of the Southern Baltic Sea: the Kiel Bight (KB, orange squares), and the Bornholm Basin (BB, blue squares) (Table S1). B) Set up and data-processing: The pico-phytoplankton fraction was obtained through size fractioning on board. Upon returning to the Institute for Marine Ecosystem and Fishery Science Hamburg (IMF), we established six technical replicates per community for each dilution across the three assay temperatures (15°C, 18°C, and 22°C). We report the slopes of three traits across the dilution steps for each region*assay temperature combination. C) Proof-of-concept via meta-barcoding and flow cytometry: To test our experimental design, we obtained MOTUs (Meta-barcoding Operational Taxonomic Units) and flow cytometric fingerprints (based on size and photopigment composition). While there were some regional differences in initial MOTU composition, those were overall not significant. (Figures S1 and S2).

## Methods

### This is a methods summary. The full methods are available in the supporting information

We obtained pico-phytoplankton community samples during two RV ALKOR cruises (AL505 and AL507 respectively) in 2018 (see Figure 1 and Table S1 for sampling dates and locations) from 5m. Samples were size fractioned to obtain the pico-phytoplankton community. To rule out effects of parameters other than temperature and diversity during the experiment, all samples were grown in f/2 media [29] at the salinity of the sampling location. Community samples grew in semi-continuous batch culture in a common garden at 18°C and 100 µmol quanta m^-2^ s^-1^ (12:12 light/dark cycle) until used for the experiment. We counted cell numbers in all samples the flow cytometre, and the flow cytometric fingerprints also allow for an estimate of phenotypic diversity or trait-level diversity [30] (based on photo-pigment composition and size [31,] Figure S2 for details). Samples were then diluted in 10-fold dilution steps at the appropriate salinity, down to the lowest point of dilution (in theory containing no more than 1 species or pico-phyoplankton per mL). Six technical replicates of each sample were left to regrow to 10^6^ cells mL^-1^ at the assay temperatures of 15°C, 18°C, and 22°C. Then, we re-diluted all samples to the same density and tracked a full growth curve. Cell size was obtained from the flow cytometre to calculate an estimate of pg carbon per mL after [32].

Net photosynthesis rates were obtained when samples were in exponential phase, on PreSens ® SDR Sensor Dish optodes at 10^5^ cells mL^-1^ in the measurement vials and measured oxygen production for 15 minutes in the light, and respiration for 15 minutes in the dark.

We obtained two measures of biodiversity in our samples. One, following CTAB DNA extractions [33], samples were sent for DNA-meta-barcoding at biome-id, resulting in a MOTU (meta-barcoding operational taxonomic units) estimate for those samples. Two, phenotypic diversity [30,31] was assessed using the parameters returned by the flow cytometre.

### Statistical analysis

All data were analysed in the R programming environment (version 3.5.3.). To analyse the shape of the growth curves, non-linear curve fitting of a baranyi growth model [34] was carried out using the ‘nlsLM’ function in the R package, ‘minpack.lm’(version 1.2-1). For multi-model selection, we computed small sample-size corrected AIC scores (AICc) and then compared the models by calculating delta AICc values and AICc weights using the “MuMIn” package (version 1.42-1).

## Results

### Proof of concept

MOTU (meta-barcoding operational taxonomic units) analyses revealed that biodiversity (as species richness) was slightly higher for samples from the Kiel Bight than from the Bornholm Basin sampling region (see Figure 1C and Figure S1), and roughly in line with previous studies on Southern Baltic Sea phytoplankton communities [35]. Further, while diversity was not reduced to a single species in the most dilute samples, MOTU data confirmed that i) the most dilute samples were on average 10^5^ times less species-rich than the least dilute samples, ii) MOTU diversity scaled with phenotypic diversity (Figure S1) and finally, that diversity once established by dilution, did not change significantly throughout the time of the growth curve (Figure S2)

### Assay temperature and biogeography explain differences in functional redundancy

We consider a result to be in line with high functional redundancy, when a function - such as biomass production or net photosynthesis - does not change significantly across levels of species richness. This results in slopes across dilution steps (i.e. species richness) that are equal or close to zero. When a function declines with species richness, we assume low functional redundancy. In these cases, the slope of trait value as a function of species richness is *positive*. We report the steepness of these slopes across three assay temperatures in Figure 2 (Table S2 for slopes, and Tables S3 to S5 for model selection and output).

**Figure 2:**
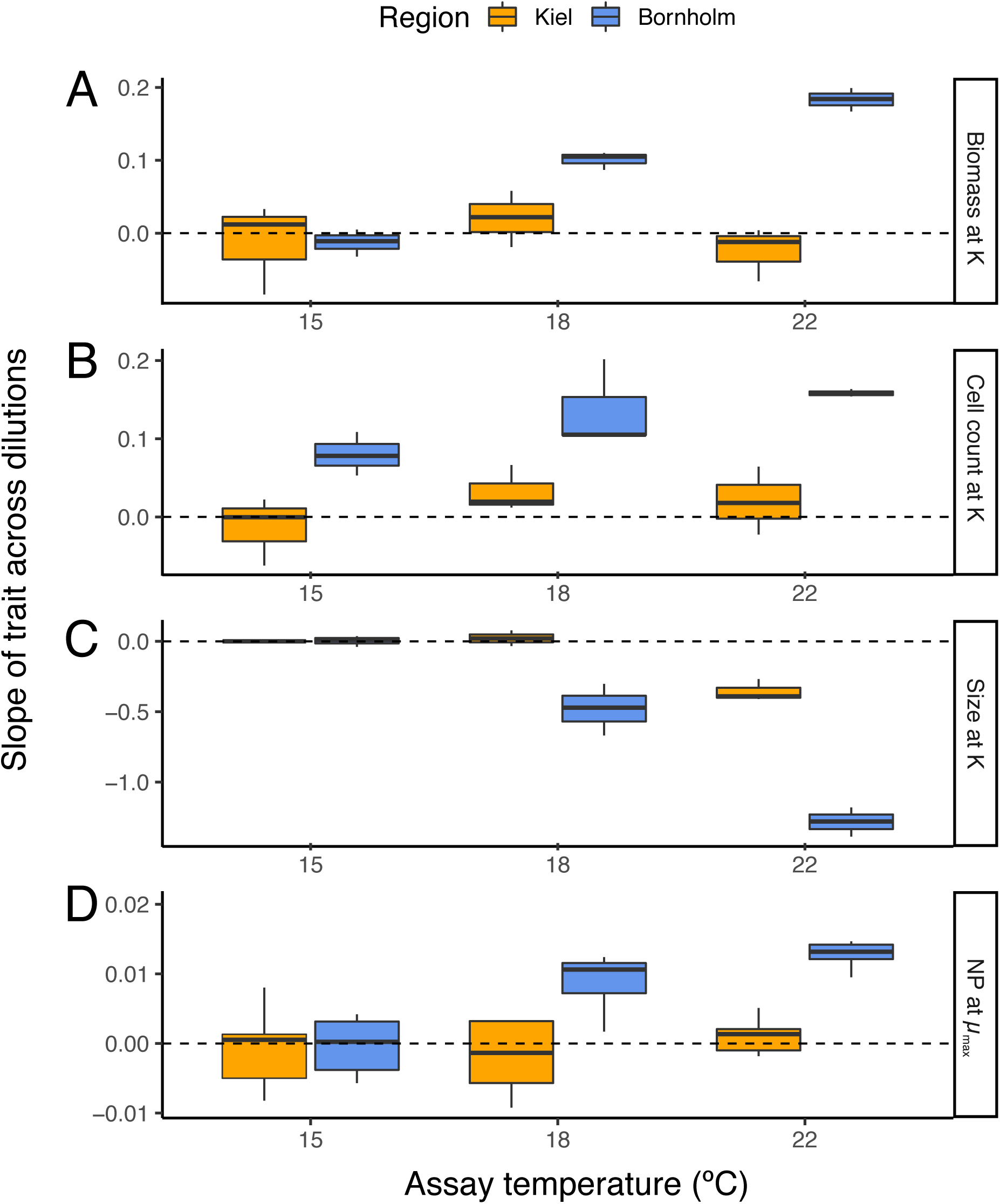
Steepness of slopes for traits across dilution gradients. (see Figures S3 to S6 and Table S2 for full slopes). The dashed line indicates a slope of 0, indicating high functional redundancy, i.e. the trait does not change significantly as rare species are lost. A positive slope indicates low functional redundancy, i.e. function decreases as species are lost. A negative slope also indicates that function changes with species richness, and specifically shows that a trait value decreases as species richness increases. **A**: Biomass produced at carrying capacity is stable across dilution steps and temperatures for samples from the Kiel Bight (high functional redundancy), however, there is low functional redundancy in samples from the Bornholm Basin, especially under warming. **B:** Cell count (mL^-1^) at carrying capacity. Cell count followed the same pattern as overall biomass. **C:** Cell size at carrying capacity. Average cell diameters were independent from dilution at 15°C for samples from either region, but cells became smaller faster with temperature in the full communities as compared to low species richness samples. This effect was more pronounced for samples from the Bornholm Basin. **D**: Net Photosynthesis (NP) across dilution steps during exponential growth showed similar patterns as the biomass response. Orange for Kiel Bight samples, blue for Bornholm Basin samples. The boxplots are displayed as is standard, with the girdle band indicating the median, and the whiskers extending to the 25th and 75th percentile. n=6 for each unique combination. This plot only displays significant parameters, i.e. samples from different seasons have been pooled.

Across dilutions, temperature had a significant impact on biomass (Figure S3, likelihood ratio test comparing models with and without ‘temperature’: Δ d.f. = 4, *χ*2 = 382.71, *P* < 0.0001, Table S3). When communities lost rare species, biomass production did not change significantly in the samples from the Kiel Bight (Figure 2A). Biomass production in samples from the Bornholm Bight samples rapidly decreased when rare species were lost, resulting in positive slopes (Figure S3, Figure 2A, likelihood ratio test comparing models with and without ‘region’: Δ d.f. =2, *χ*2 = 185.23, *P* < 0.0001, Table S3).

Changes in biomass were driven by a change in cell number and a change in cell volume across dilution steps, with a trend for smaller cells at higher species richness (Figure 2B, C, Figure S4, S5, Table S2, S4). Cell volume decreased with temperature (likelihood ratio test comparing models with and without ‘temperature’: Δd.f. = 2, *χ*2 =382.70, *P* < 0.0001, Figure 2C, Table S4). While cells in samples from the cooler, more stable, Bornholm Basin were on average less reactive to temperature in terms of cell size, the effect of losing rare species was stronger here than in samples from the Kiel Bight and cell size rapidly increased as species richness decreased (likelihood ratio test comparing models with and without ‘region: Δd.f. = 2, *χ*2 = 223.68, *P* < 0.0001, Figure 2 C, Table S4)

Net Photosynthesis rates (NP) per cell followed a similar trend to biomass (Figure 2D, Figure S6, Table S5 for model selection and output): assay temperature altered community NP depending on dilution and sampling origin. Samples from the Kiel region were overall more photosynthetically active than those from the Bornholm region (likelihood ratio test comparing models with and without ‘region’: Δ d.f. = 3, *χ*2 = 21.63, *P* < 0.0001, Figure 2D, Figure S6, Table S5), especially at the warmer temperatures (likelihood ratio test comparing models with and without ‘temperature’: Δ d.f. = 1, *χ*2 = 85.47, *P* < 0.0001, Figure 2D Figure S6, Table S5). NP in samples from the Kiel region showed no clear trend with dilution. In samples from the Bornholm region, on the other hand, responses of NP to temperature were overall less pronounced, but were strongly influenced by species richness, with the full community samples photosynthesising nearly 1.5 times (per cell) as much as the samples with the lowest species richness.

## Discussion

Our investigations of the link between species richness and traits relevant for ecosystem services showed that the steepness of the biodiversity-function slope hinged on the communities’ evolutionary histories as well as on short-term (ca 20 generations) changes in temperature and fit well within the theoretical and observational general framework of biodiversity/ecosystem function studies [2-5]. The effect of temperature modulating the strength of the diversity/function relationship was the most pronounced for samples originating from a cooler, less variable region, in line with theory predicting that regions that are more variable should contain a greater number of taxa with more variable tolerance thresholds [36]. Overall, our findings are also in accordance with the literature on experimentally assembled communities in microcosms, where the researchers found that a decline in productivity was more pronounced at rising temperatures in phytoplankton and bacteria ([10] and [9] respectively). We make the case that this pattern of declining productivity due to species loss being enhanced at higher temperatures may be conserved across environments as diverse as laboratory conditions and natural community samples. This may also mean that results obtained from assembling artificial communities reflect results found in natural assemblages well in direction, and possibly in magnitudes of responses. We specifically find that as species are lost from samples with a history of comparatively cooler average temperatures and highly predictable seasonal variation, the community rapidly shows a decrease in net photosynthesis rates, biomass production, and – unlike in experimental assemblages [10] - an increase in cell size. Changes in the biodiversity–ecosystem function relationship were thus strongly linked to size-dependent turnover in community composition, but it is unclear whether this is a cause or a consequence of the other observed patterns. We can only speculate about the underlying mechanisms. While it is possible that competition for resources is stronger in a mixed community, and that this pressure is relieved once most species are lost from the system, our samples were kept in full media, and the pattern was obvious during exponential phase already (Figure S7). It is thus unlikely that competition for nutrients, light, or space, was the main factor here. Neither can the pattern be explained by the larger cells’ being more common in the original samples (which do have slightly higher variation in cell size than very dilute samples) and therefore more likely to remain present after dilution. The pattern may be driven by interactions beyond competition, e.g. types of facilitation [37], or a more general trend for trait variability and biodiversity being intertwined [38]. It remains to be seen to which degree the patterns we observed here are also driven by interactions with their associated bacteria. The latter may have different nutrient requirements that are not met in the same way by our culture media, which may in turn affect how they interact with and shape the phenotypes of the phytoplankton species.

Irrespective of the reasons underlying the shift in cell size we observed here, the contribution of the very small pico-phytoplankton cells to total phytoplankton biomass in the ocean has been shown to increase with temperature [39] and changes to size distribution in scenarios where warming and loss of biodiversity interact may have unpredictable impacts on the function of phytoplankton ecosystems [40]

While biodiversity manipulation by dilution has limitations (e.g. diversity and species identity can never be fully disentangled, dilution introduces quasi-random differences in beta diversity), these apparent disadvantages have potential important ecological implications as they allow us to specifically investigate the importance of rare taxa, rather than random species loss.

The Baltic Sea is generally low in species due to its characteristic brackish waters. To rule out that our results are typical for regions where biodiversity (as richness) is low to begin with, it would be necessary to carry out comparable studies across systems that vary in their initial species compositions. Together, such data will further improve our understanding of the relationship between diversity and ecosystem function at the foundation of warming aquatic ecosystems.

## Supporting information

Supporting Images and Tables

## Author contributions

DZ carried out the experiments. LL and MS retrieved, prepared, and maintained the phytoplankton cultures. ES conceived the experiment, supervised laboratory work and handled data analysis. All authors contributed equally to writing the manuscript.

## Acknowledgements

We would like to thank Margarethe Nowicki, Richard Klinger, Jens-Peter Hermann, and the captain and crew of RV ALKOR for support at sea (cruises AL505 and AL507), and Stefanie Schnell for technical assistance in the laboratory at IMF Hamburg. Samuel Barton, Moritz Aehle, and Paula Franze assisted with the metabolism measurements.

