## Supporting Images and Tables for "Functional redundancy in natural pico-phytoplankton communities depends on temperature and biogeography"

Author affiliations:
[1] University of Hamburg, Institute for Marine Ecosystem and Fisheries Science, 22767 Hamburg
[2] Centre for Earth System Science and Sustainability, 20146 Hamburg

Corresponding author


### Full methods

We obtained pico-phytoplankton community samples during two RV ALKOR cruises (AL505 and AL507 respectively) in 2018 (see Figure 1 and Table S1 for sampling dates and locations) using a Niskin bottle at 5m. Samples were immediately passed through a 35µm sieve to remove grazers and large debris, and then further size fractioned *via* gentle filtration through a 2µm membrane filter (kept filtrate) and an 0.2µm filter (kept filter and rinsed gently). Acute thermal profiles for the communities were determined during on-board incubations (Santelia *et al.*, in prep. – SI document 2) in order to better be able to estimate which temperatures to use as assay temperatures in the laboratory. To rule out effects of parameters other than temperature and diversity during the experiment, all samples were grown in f/2 media [34] at the salinity of the sampling location. Community samples grew in semi-continuous batch culture in a common garden at 18°C and 100 µmol quanta $m^{-2}$ $s^{-1}$ (12:12 light/dark cycle) until used for the experiment. To set up the experiment, we first counted cell numbers in three representative Kiel Bight community samples (i.e. 3 stations) and three representative Bornholm Basin samples (i.e. 3 stations) using a BD Accuri C6 flow cytometre. The flow cytometric fingerprints also allow for an estimate of phenotypic diversity or trait-level diversity [35] which is largely based on photo-pigment composition and size [36] (see also Figure S2 for details on analysis). Samples were then diluted in 10-fold dilution steps at the appropriate salinity, down to the lowest point of dilution (in theory containing no more than 1 species or pico-phyoplanktonper mL). Six technical replicates of each sample were left to regrow to $10^6$ cells $mL^{-1}$ at the assay temperatures of 15ºC, 18ºC, and 22ºC. These temperatures are all within the ranges of temperatures commonly experienced during late spring (15ºC), summer (18ºC), and the height of summer (22ºC). This resulted in a total of 270 experimental units. Then, we re-diluted all samples to 3000 cells $mL^{-1}$ and tracked a full growth curve until samples reached carrying capacity (ca. 23 days) in all experimental units at all temperatures, with measurements taken on the flow cytometre every other day. Cell size as diameter in μm was obtained from the flow cytometre's forward scatter after calibration with size beads. Taking into account cell counts per mL and assuming on average spherical shapes and using conversion factors after [37], we then calculated an estimate of pg carbon per mL to obtain biomass produced.

Net photosynthesis rates were obtained when samples were in exponential phase, on PreSens ® SDR Sensor Dish optodes. Here, we aimed for a total of $10^5$ cells $mL^{-1}$ in the measurement vials and measured oxygen production for 15 minutes in the light, and respiration for 15 minutes in the dark. Net photosynthesis was calculated considering that phytoplankton in our set-up will only be able to photosynthesise during the light phase (12 hours), but will respire throughout the day *and* night phase (24 hours). All measurements were carried out at the same time of day ( ~ 9am to 11am) under the light- and temperature conditions set in the incubator (i.e. all experimental units at their assay temperatures).

We obtained two measures of biodiversity in our samples. One, following CTAB DNA extractions [38], a subset of representative samples was sent for DNA- meta-barcoding at biome-id (16S primers: forward CCTACGGGNGGCWGCAG, and reverse GACTACHVGGGTATCTAATCC, 18S primers: forward CCGCGGTAATTCCAGCTC and reverse CCTTGGTCCGTGTTTCTAGAC), resulting in a MOTU (meta-barcoding operational taxonomic units) estimate for those samples. Two, phenotypic diversity [35,36] was assessed using the parameters returned by the flow cytometre.

**Statistical analysis**

All data were analysed in the R programming environment (version 3.5.3.). To analyse the shape of the growth curves, non-linear curve fitting of a baranyi growth model [39] was carried out using the 'nlsLM' function in the R package, 'minpack.lm'(version 1.2-1).

Parameter estimation was achieved by running 1,000 different random combinations of starting parameters for cell count at carrying capacity, duration of lag phase, and maximum growth rate picked from a uniform distribution. The script then retains the parameter set that returned the lowest Akaike information criterion (AICc) score. Parameters (biomass and cell size at carrying capacity, net photosynthesis during exponential growth) were then compared through a mixed effects model (within the nlme package, version 3.1-137). There, the respective parameters were explained by a global model that included sampling location (Kiel Bight or Bornholm Basin), assay temperature (15ºC,18ºC, or 22ºC), and dilution step (from lowest to highest) and sampling season (spring or summer) as fixed factors in full interaction. Sampling station was computed as a nested random effect within region. In all cases, seasonality was found to not explain the data better and was subsequently dropped from the fixed factors to avoid over-parameterisation of the model. For multi-model selection, we computed small sample-size corrected AIC scores (AICc) and then compared the models by calculating delta AICc values and AICc weights using the "MuMIn" package (version 1.42-1). We picked the model where delta AICc was > 2 for refitting with REML. For graphical presentation of data, we used the ggplot2 (version 3.2) and vegan (ordihull and metaMDS, version 2.5-4, see SI for details on NMDS plots) packages.

### Supporting Figures

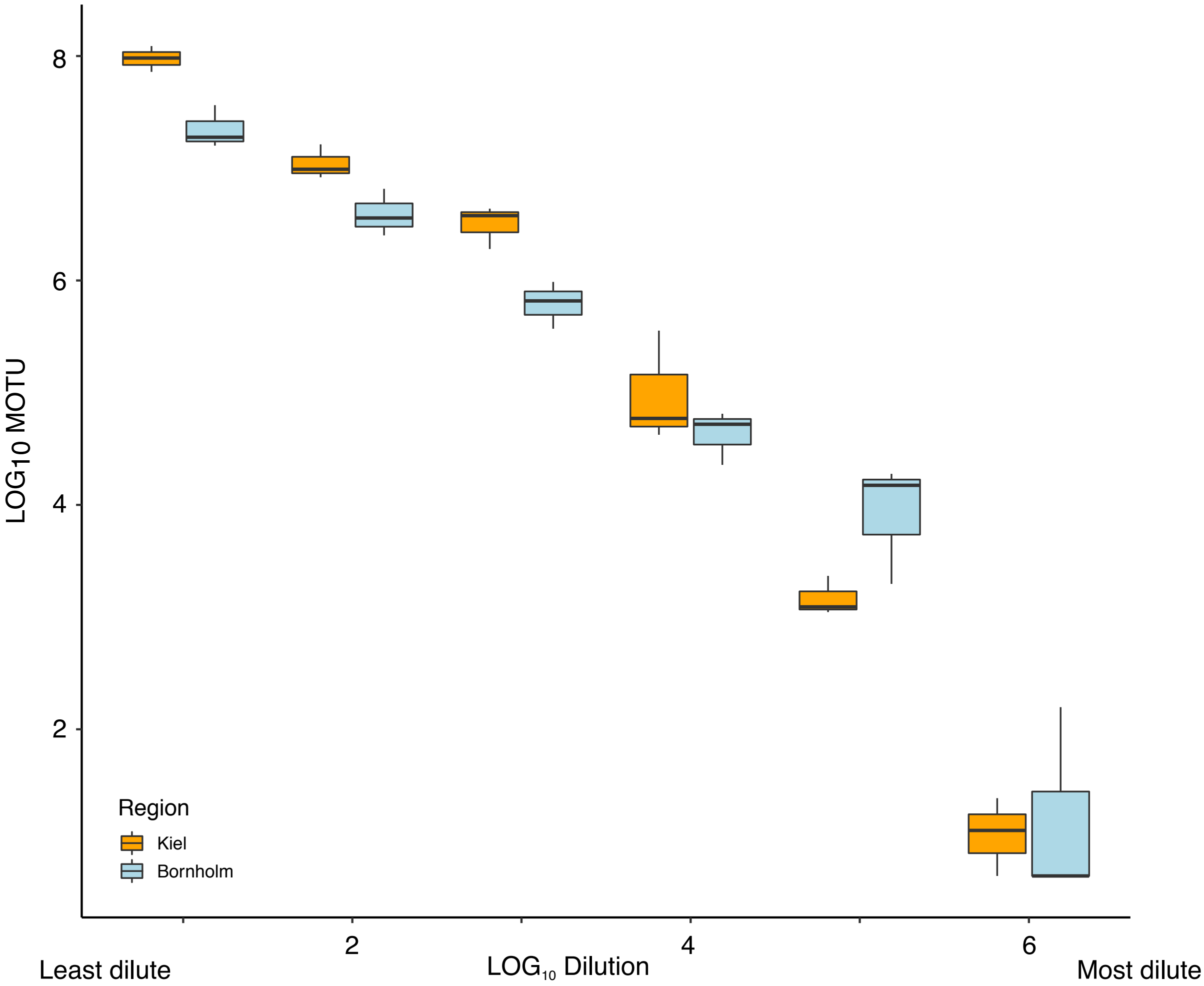

**Figure S1: Dilution is strongly correlated to MOTU (operational taxonomic units obtained through meta barcoding):**

The relationship between the logarithms of dilution and MOTU count reveals that the dilutions successfully reduced species richness in Kiel Bight (orange) and Bornholm Basin (blue) samples. Kiel samples had slightly higher original MOTU counts, which was driven largely by a slightly lower species count and higher predominance of cyanobacteria in the Bornholm Basin samples during the summer (see also below). The boxplots are displayed as is standard, with the girdle band indicating the median, and the whiskers extending to the 25th and 75th percentile. For each unique treatment combination (dilution*region*temperature), n=6.

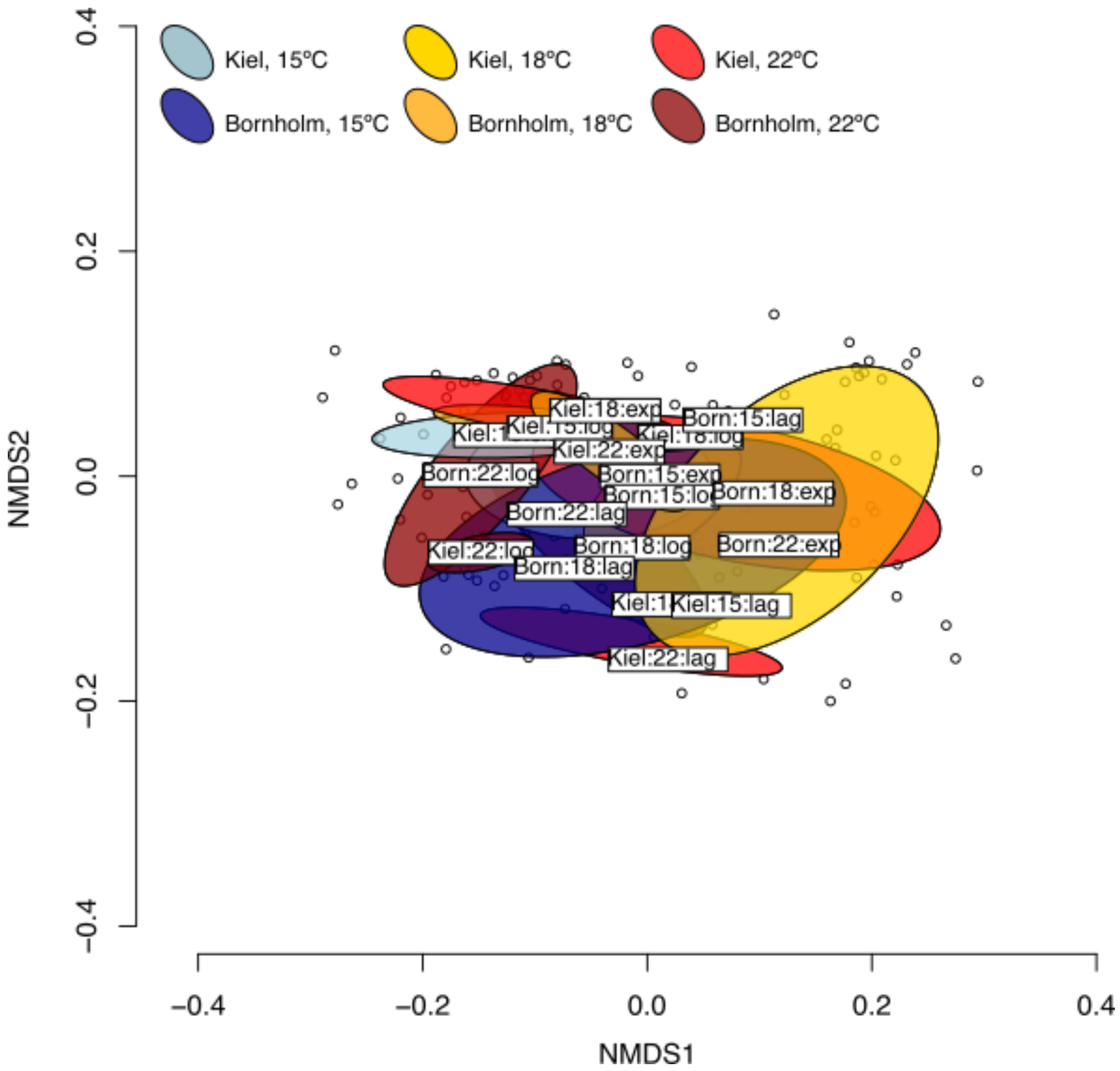

**Figure S2: Once established through dilution, community phenotypic diversity remained largely stable throughout the growth cycle.**

Variation in the composition of the phytoplankton communities between treatments (15ºC in blue tones, 18ºC in orange and yellow, 22ºC in red tones), time points (lag for lag phase, exp for exponential phase, and log for time of carrying capacity), and among sampling regions (lighter hues for Kiel Bight and darker hues for Bormholm Basin) was indexed as the "score" for each unique combination of sampling region, temperature, and time point along the first axis of a non-metric multidimensional scaling (NMDS) ordination. Composition of phytoplankton communities was investigated *via* flow cytometric fingerprints, i.e. taking into account size (FSC), granularity (SSC), relative chlorophyll a content (FL3), relative phycoeryhtrin content (FL2), and relative allophycocyanine content (FL4) (see e.g [1,2]). NMDS ordination was conducted using the "metaMDS" function in the "vegan" package in R based on a Jaccard dissimilarity matrix. We used Permutational Multivariate Analysis of Variance (PERMANOVA) to test whether dissimilarities between temperatures, dilutions, time points and temperatures or their interaction caused significantly different population makeups. Samples from the Bornholm region had a borderline significantly different make up of species ($F_{1,13}$= 2.72, p =0.058) driven solely by the summer months seeing a higher abundance of cyanobacteria in the Bornholm region, (see also Santelia *et al.*, in prep.- SI document 2), but there was no significant change in community composition throughout the growth curve for individual dilution steps ($F_{2,13}$= 2.35, p =0.08).

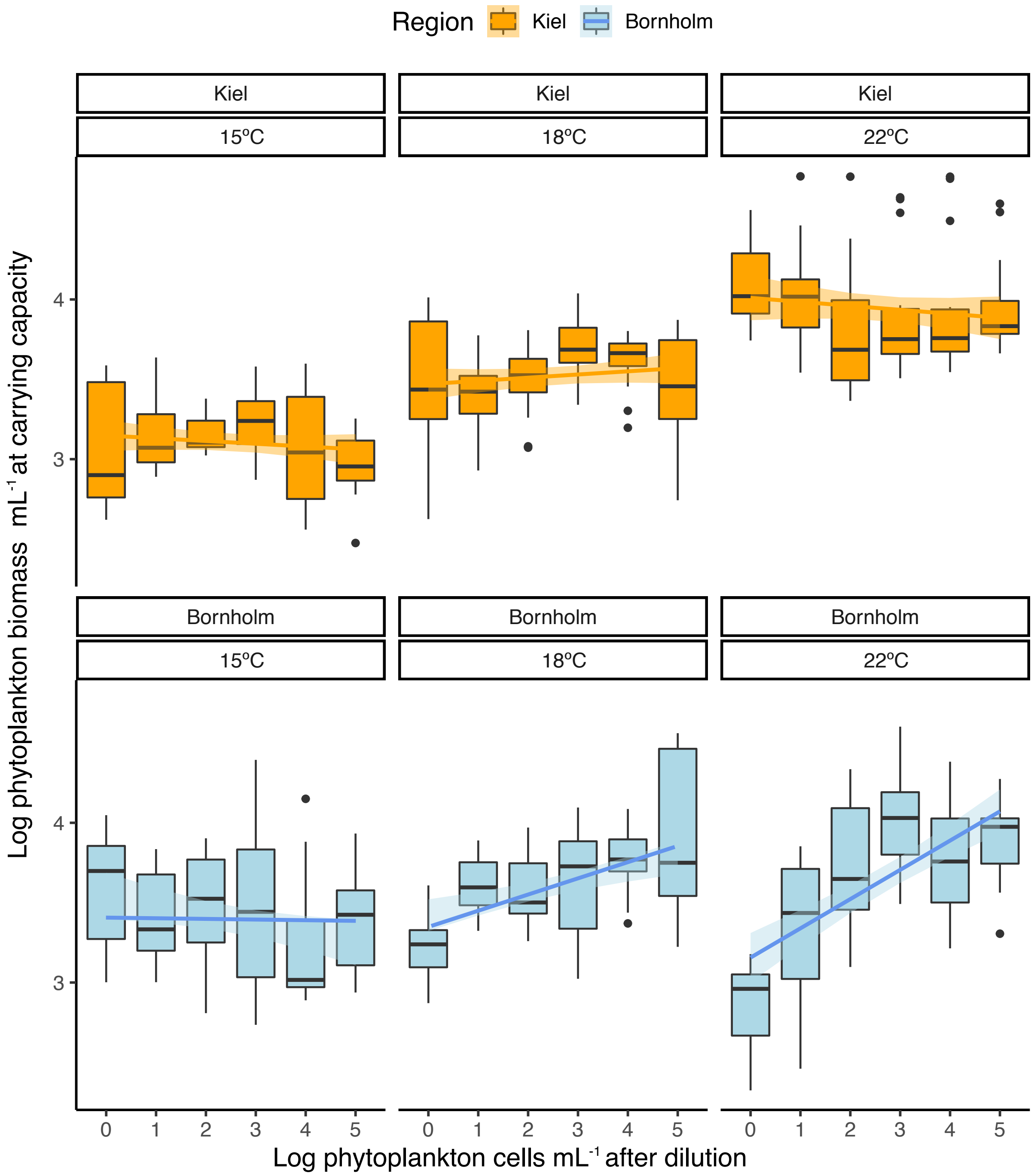

**Figure S3: Biomass at carrying capacity K**

Biomass at carrying capacity (here in pg C per mL, displayed as LOG10 for clarity) in samples from the Kiel sampling stations (orange, upper) and the Bornholm sampling stations (blue, lower) was influenced by assay temperature (individual panels) and dilution (here, displayed as the LOG10 of phytoplankton cells after dilution, which is a good indicator for species richness (see Figure S1 and main manuscript Figure 1). A slope that does not deviate significantly from 0 (see also Table S2) indicates that functional redundancy is high. A slope that does deviate significantly from 0 indicates that species richness has a strong impact on the trait under investigation, with positive slopes for samples with low functional redundancy. While temperature has an impact on biomass at carrying capacity rates in the samples from the Kiel Area (highest rates at 18ºC, lowest at 15ºC, and intermediate values for 22ºC), there is no significant impact of loss of rare species (i.e. dilution). The boxplots are displayed as is standard, with the girdle band indicating the median, and the whiskers extending to the 25th and 75th percentile. For each unique treatment combination (dilution*region*temperature), n=6.

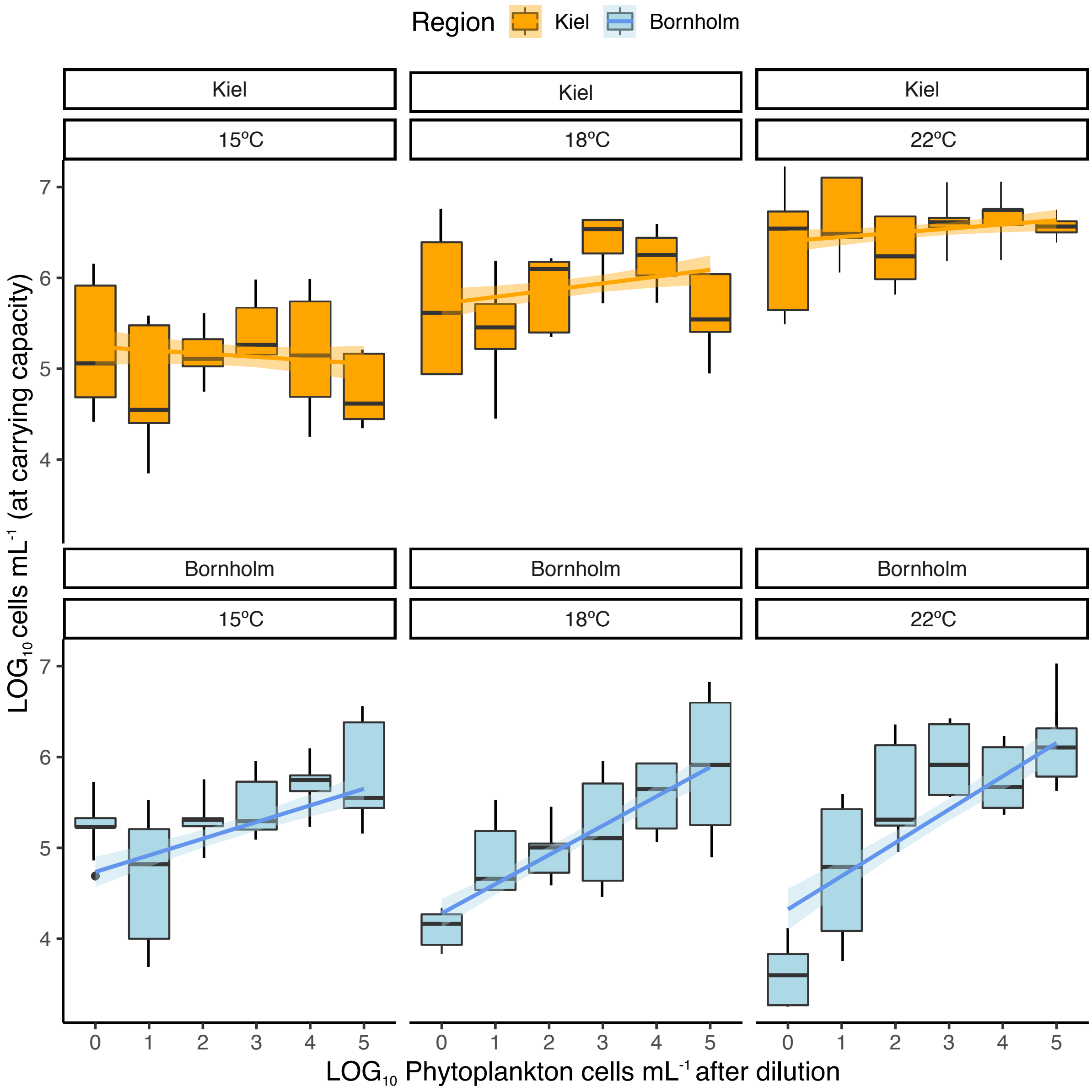

**Figure S4: Cell count at carrying capacity**

Cell count mL$^{-1}$ at carrying capacity (here displayed as LOG10 for clarity) in samples from the Kiel sampling stations (orange, upper) and the Bornholm sampling stations (blue, lower) was influenced by assay temperature (individual panels) and dilution (here, displayed as the LOG10 of phytoplankton cells after dilution, which is a good indicator for species richness (see Figure S1 and main manuscript Figure 1). A slope that does not deviate significantly from 0 (see also Table S2) indicates that functional redundancy is high. A slope that does deviate significantly from 0 indicates that species richness has a strong impact on the trait under investigation, with positive slopes for samples with low functional redundancy. While temperature has an impact on cell count at carrying capacity rates in the samples from the Kiel Area (highest rates at 18ºC, lowest at 15ºC, and intermediate values for 22ºC), there is no significant impact of loss of rare species (i.e. dilution). The boxplots are displayed as is standard, with the girdle band indicating the median, and the whiskers extending to the 25th and 75th percentile. For each unique treatment combination (dilution*region*temperature), n=6.

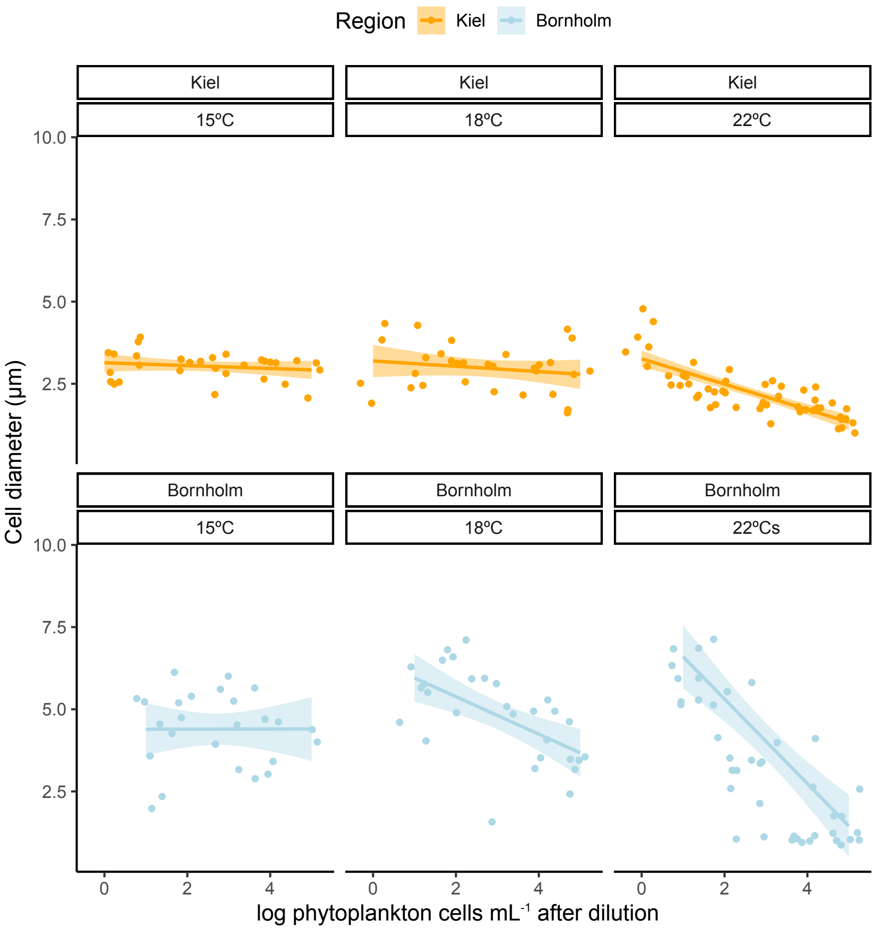

**Figure S5: Size (diameter in µm) at carrying capacity as used for biomass estimation**
At carrying capacity, cell diameter in samples from the Kiel sampling stations (orange, upper) and the Bornholm sampling stations (blue, lower) was influenced by assay temperature (individual panels) and dilution (here, displayed as the LOG10 of phytoplankton cells after dilution, which is a good indicator for species richness (see Figure S1 and main manuscript Figure 1). At 15ºC, dilution did not significantly affect cell size. At 18ºC, dilution affected cell size only in samples from the Bornholm region. At the highest temperature (22ºC), cell size strongly decreased when communities were more diverse in samples from both regions. A slope that does not deviate significantly from 0 (see also Table S2) indicates that functional redundancy is high. A slope that does deviate significantly from 0 indicates that species richness has a strong impact on the trait under investigation. For each unique treatment combination (dilution*region*temperature), n=6.

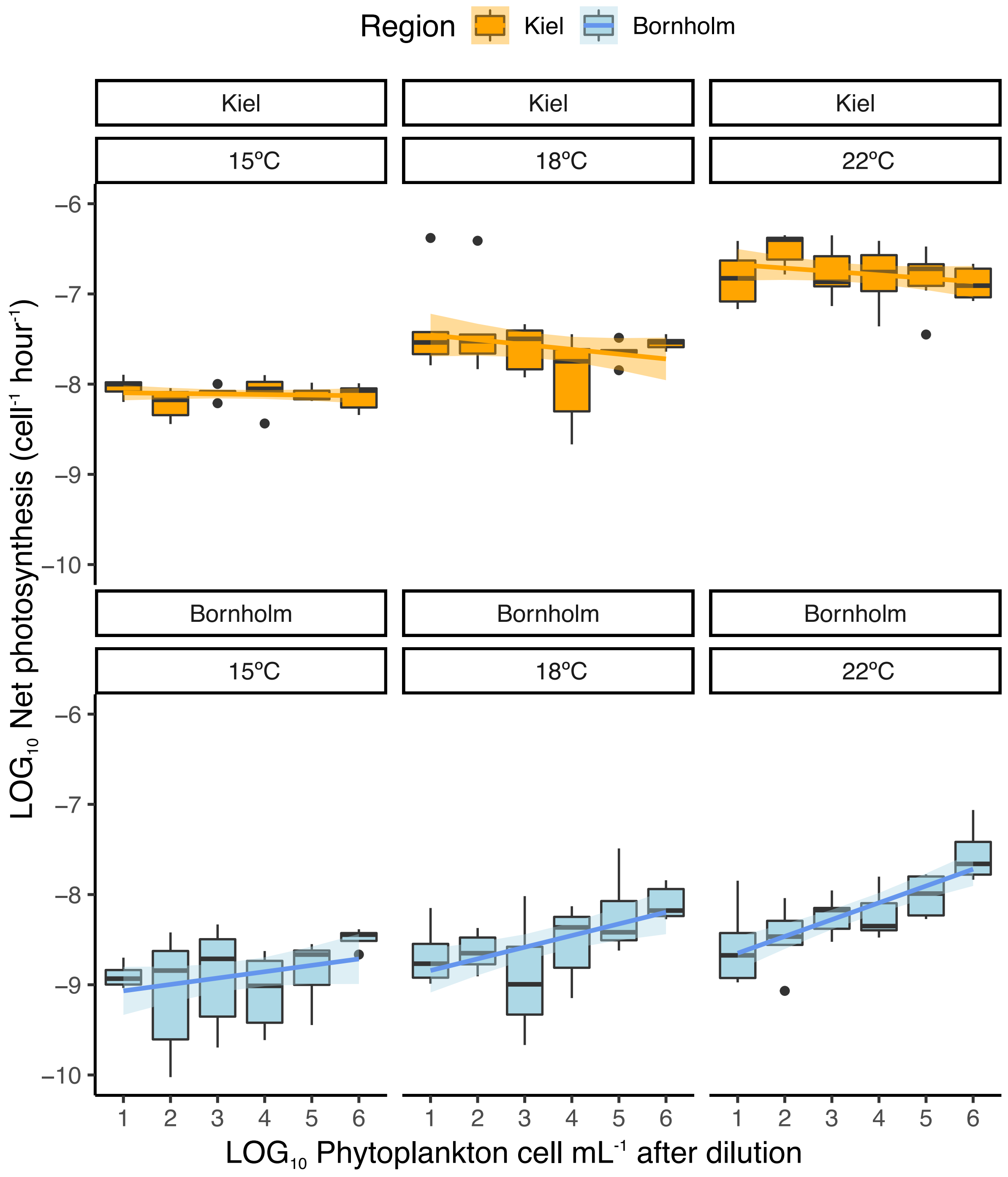

**Figure S6 Rates of net photosynthesis (**µmol $O_2$ per cell and hour**) during exponential growth**

During exponential growth, Net Photosynthesis (NP, here in µmol $O_2$ per cell and hour) in samples from the Kiel sampling stations (orange, upper) and the Bornholm sampling stations (blue, lower) was influenced by assay temperature (individual panels) and dilution (here, displayed as the LOG10 of phytoplankton cells after dilution, which is a good indicator for species richness (see Figure S1 and main manuscript Figure 1). A slope that does not deviate significantly from 0 (see also Table S2) indicates that functional redundancy is high. A slope that does deviate significantly from 0 indicates that species richness has a strong impact on the trait under investigation, with positive slopes for samples with low functional redundancy. While temperature has an impact on gross photosynthesis rates in the samples from the Kiel Area (highest rates at 18ºC, lowest at 15ºC, and intermediate values for 22ºC), there is no significant impact of loss of rare species (i.e. dilution). In samples from the Bornholm Basin, NP  rates are overall lower, and samples with lower species richness are significantly less photosynthetically active than samples with high species richness, and this trend is exacerbated with increasing temperatures. For each unique treatment combination (dilution*region*temperature), n=6. The boxplots are displayed as is standard, with the girdle band indicating the median, and the whiskers extending to the 25th and 75th percent

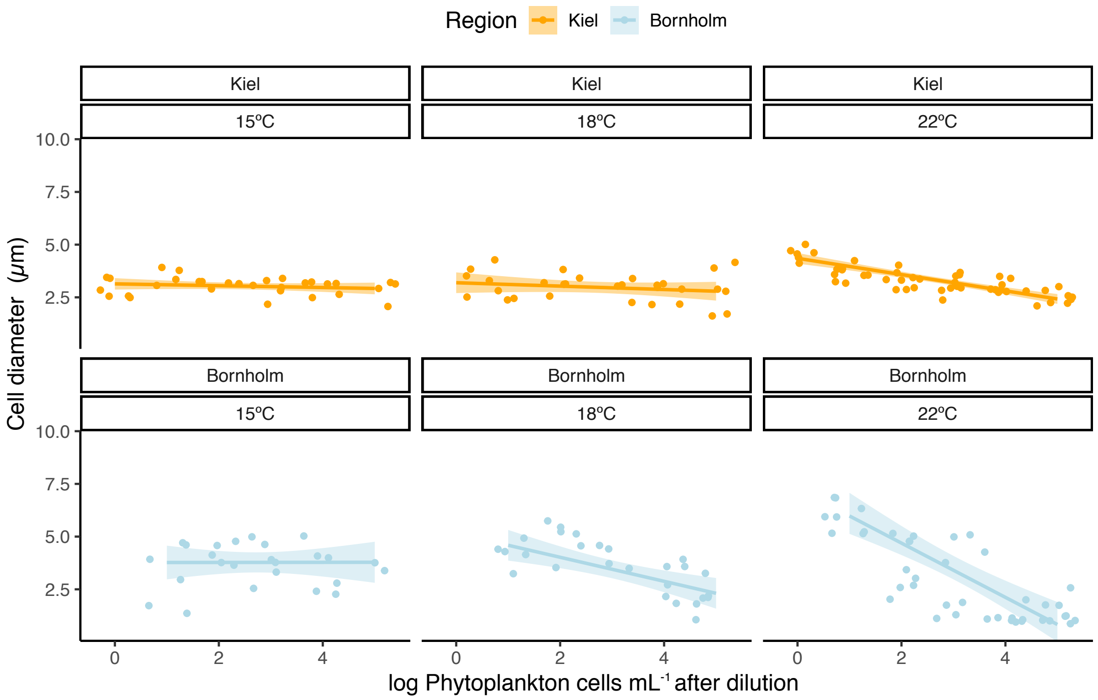

**Figure S7 Size (diameter in µm) during exponential growth**

During exponential growth, cell diameter in samples from the Kiel sampling stations (orange, upper) and the Bornholm sampling stations (blue, lower) was influenced by assay temperature (individual panels) and dilution (here, displayed as the LOG10 of phytoplankton cells after dilution, which is a good indicator for species richness (see Figure S1 and main manuscript Figure 1). The pattern observed at carrying capacity (see Figure S5) is also reflected during exponential growth. A slope that does not deviate significantly from 0 (see also Table S2) indicates that functional redundancy is high. A slope that does deviate significantly from 0 indicates that species richness has a strong impact on the trait under investigation. For each unique treatment combination (dilution*region*temperature), n=6.

### Supporting Tables

##### Table S1: Sampling station overview

Below, we provide the coordinates (Long/Lat), time of sampling (spring or summer 2018 and official ALKOR identifier), as well as salinity, temperature, and nutrient content at time of sampling for each station. See also map in Figure 1 in the main text. Three technical replicates were established for each Station at each temperature and each level of dilution. StationID is as used throughout this manuscript, and not an official station identifier. As each individual station was only sampled once (with two technical replicates) for nutrient content, temperature, and salinity as is standard, we do not provide standard deviations as they would not carry any true meaning. Temperature and salinity data are as exported from the ship's CTD. Nutrient content was measured on a SEAL sequential analyser (AA3) following protocols of [3,4] upon returning to Hamburg.

| StationID | Time of sampling /cruise ID | Longitude | Latitude | Temperature (ºC) | Salinity | Nitrate+ nitrite (µg mL$^{-1}$) | Phosphate (µg mL$^{-1}$) | Silicate (µmol L$^{-1}$) |
|---|---|---|---|---|---|---|---|---|
| Kiel01 | Spring 2018 (AL505) | 11º19.36 | 54º31.27 | 1.76 | 11.16 | 53.69 | 18.97 | 12.56 |
| Kiel02 | Spring 2018 (AL505) | 10º40.51 | 54º42.42 | 1.28 | 13.15 | 51.58 | 16.29 | 13.12 |
| Kiel03 | Summer 2018 (AL507) | 10º20.22 | 54º41.7 | 21.35 | 15.00 | 21.17 | 4.56 | 5.44 |
| Bornholm01 | Spring 2018 (AL505) | 15º54.06 | 55º44.16 | 2.03 | 7.36 | 44.21 | 20.08 | 10.47 |
| Bornholm02 | Spring 2018 (AL505) | 15º26.02 | 55º16.87 | 2.43 | 7.44 | 46.64 | 22.17 | 12.88 |
| Bornholm03 | Summer 2018 (AL507) | 15º08.87 | 54º53.08 | 22.6 | 6.5 | 15.53 | 2.22 | 4.87 |

**Table S2: Slopes obtained (per station) from Figures S3 to S5**

Slopes for each temperature, region, and station for each trait investigated. Technical replicates have been pooled (to statID), and the slope reported is the average, with 1SD in 'sd_slope'. NP is for net photosynthesis. NP has been established during exponential growth. Biomass, cell count, and cell size were established during carrying capacity.

| Temp | Region | statID | slope | Trait | sd_slope |
|---|---|---|---|---|---|
| 15 | Kiel | 1 | 0.012 | Biomass | 0.007 |
| 15 | Kiel | 2 | 0.033 | Biomass | 0.019 |
| 15 | Kiel | 3 | -0.084 | Biomass | 0.049 |
| 15 | Bornholm | 1 | 0.005 | Biomass | 0.003 |
| 15 | Bornholm | 2 | -0.032 | Biomass | 0.019 |
| 15 | Bornholm | 3 | -0.011 | Biomass | 0.006 |
| 18 | Kiel | 1 | 0.022 | Biomass | 0.013 |
| 18 | Kiel | 2 | 0.058 | Biomass | 0.033 |
| 18 | Kiel | 3 | -0.019 | Biomass | 0.011 |
| 18 | Bornholm | 1 | 0.087 | Biomass | 0.05 |
| 18 | Bornholm | 2 | 0.105 | Biomass | 0.061 |
| 18 | Bornholm | 3 | 0.11 | Biomass | 0.064 |
| 22 | Kiel | 1 | 0.004 | Biomass | 0.002 |
| 22 | Kiel | 2 | -0.066 | Biomass | 0.038 |
| 22 | Kiel | 3 | -0.012 | Biomass | 0.007 |
| 22 | Bornholm | 1 | 0.167 | Biomass | 0.096 |
| 22 | Bornholm | 2 | 0.199 | Biomass | 0.001 |
| 22 | Bornholm | 3 | 0.184 | Biomass | 0.106 |
| 15 | Kiel | 1 | 0 | Cell count | 0.031 |
| 15 | Kiel | 2 | 0.022 | Cell count | 0.025 |
| 15 | Kiel | 3 | -0.062 | Cell count | 0.022 |
| 15 | Bornholm | 1 | 0.053 | Cell count | 0.02 |
| 15 | Bornholm | 2 | 0.108 | Cell count | 0.018 |
| 15 | Bornholm | 3 | 0.078 | Cell count | 0.018 |
| 18 | Kiel | 1 | 0.019 | Cell count | 0.021 |
| 18 | Kiel | 2 | 0.066 | Cell count | 0.017 |
| 18 | Kiel | 3 | 0.012 | Cell count | 0.029 |
| 18 | Bornholm | 1 | 0.101 | Cell count | 0.013 |
| 18 | Bornholm | 2 | 0.102 | Cell count | 0.024 |
| 18 | Bornholm | 3 | 0.196 | Cell count | 0.029 |
| 22 | Kiel | 1 | -0.022 | Cell count | 0.056 |
| 22 | Kiel | 2 | 0.064 | Cell count | 0.039 |
| 22 | Kiel | 3 | 0.017 | Cell count | 0.033 |
| 22 | Bornholm | 1 | 0.159 | Cell count | 0.008 |
| 22 | Bornholm | 2 | 0.161 | Cell count | 0.021 |
| 22 | Bornholm | 3 | 0.167 | Cell count | 0.046 |
| 15 | Kiel | 1 | -0.005 | NP | 0.009 |

| | | | | | |
|---|---|---|---|---|---|
| 15 | Kiel | 2 | -0.031 | NP | -0.003 |
| 15 | Kiel | 3 | -0.001 | NP | 0.002 |
| 15 | Bornholm | 1 | 0.076 | NP | -0.002 |
| 15 | Bornholm | 2 | -0.059 | NP | -0.002 |
| 15 | Bornholm | 3 | 0.147 | NP | 0 |
| 18 | Kiel | 1 | -0.025 | NP | 0.012 |
| 18 | Kiel | 2 | 0.007 | NP | -0.002 |
| 18 | Kiel | 3 | -0.003 | NP | -0.002 |
| 18 | Bornholm | 1 | 0.137 | NP | 0.003 |
| 18 | Bornholm | 2 | 0.121 | NP | 0.006 |
| 18 | Bornholm | 3 | 0.179 | NP | -0.001 |
| 22 | Kiel | 1 | -0.001 | NP | 0.002 |
| 22 | Kiel | 2 | 0.002 | NP | -0.001 |
| 22 | Kiel | 3 | -0.009 | NP | 0 |
| 22 | Bornholm | 1 | 0.199 | NP | 0.005 |
| 22 | Bornholm | 2 | 0.179 | NP | 0.007 |
| 22 | Bornholm | 3 | 0.181 | NP | 0.008 |
| 15 | Kiel | 1 | 0.001 | Size | 0 |
| 15 | Kiel | 2 | 0.001 | Size | 0 |
| 15 | Kiel | 3 | 0.002 | Size | 0 |
| 15 | Bornholm | 1 | 0.037 | Size | 0.021 |
| 15 | Bornholm | 2 | -0.039 | Size | 0.023 |
| 15 | Bornholm | 3 | 0.01 | Size | 0.006 |
| 18 | Kiel | 1 | 0.04 | Size | 0.015 |
| 18 | Kiel | 2 | 0.004 | Size | 0.014 |
| 18 | Kiel | 3 | 0.006 | Size | 0.014 |
| 18 | Bornholm | 1 | -0.668 | Size | 0.004 |
| 18 | Bornholm | 2 | -0.471 | Size | 0.003 |
| 18 | Bornholm | 3 | -0.302 | Size | 0.001 |
| 22 | Kiel | 1 | -0.222 | Size | 0.003 |
| 22 | Kiel | 2 | -0.371 | Size | 0.002 |
| 22 | Kiel | 3 | -0.382 | Size | 0.002 |
| 22 | Bornholm | 1 | -1.387 | Size | 0.013 |
| 22 | Bornholm | 2 | -1.18 | Size | 0.029 |
| 22 | Bornholm | 3 | -1.12 | Size | 0.018 |

**Table S3: Model selection (A) output (B) for investigating the effect of dilution (abbreviated to D), assay temperature (abbreviated T), region (abbreviated R), and season (abbreviated S) on biomass produced at carrying capacity.**

In the mixed model, D (from 1 – highest richness to 1e-05 – lowest richness), T (15ºC,18ºC, 22ºC), R (Kiel Area, Bornholm Basin), and S (spring, summer) selection regimes, i.e. nutrient (low nutrient and replete), were fitted as fixed effects. Stations were treated as a random factor. Technical replicates were not fitted. The best model is highlighted in bold, and is the model with the smallest AICc, where delta AICc to the next best model is >2. df for degrees of freedom; logLik for log likelihood ratio. : indicates an interaction term. We display only the first 10 models for clarity.

The global model formula was lme.formula(K~D*R*S*T, random=~1|bio.stat.id, data=dataframe.K, method=”ML) . The model used for the model output table was refitted with REML and read lme.formula(K~D*R*T, random=~1|bio.stat.id, data=dataframe.K, method=”REML). In the model output table, CI are the 95% confidence intervals, DF are degrees of freedom. Values other than the first value (Kiel sample at 15ºC with the lowest dilution, i.e. highest diversity) need to be added to the first value to obtain the predicted trait value.

| **A Intercept** | **D** | **T** | **R** | **S** | **D:T** | **D:R** | **D:S** | **T:R** | **S:T** | **R:S** | **D:T:R** | **D:T:S** | **D:R:S** | **T:R:S** | **D:T:R:S** | **df** | **logLik** | **AICc** | **Δ AICc** | **weight** |
|---|---|---|---|---|---|---|---|---|---|---|---|---|---|---|---|---|---|---|---|---|
| 3.06 | + | + | + | NA | + | + | NA | + | NA | NA | + | NA | NA | NA | NA | 38 | -129.2 | 339.93 | 0.00 | 0.68 |
| 3.06 | + | + | + | + | + | + | NA | + | NA | NA | + | NA | NA | NA | NA | 39 | -129.2 | 342.23 | 2.30 | 0.22 |
| 3.06 | + | + | + | + | + | + | NA | + | NA | + | + | NA | NA | NA | NA | 40 | -129.2 | 344.53 | 4.60 | 0.07 |
| 3.06 | + | + | + | + | + | + | NA | + | + | NA | + | NA | NA | NA | NA | 41 | -129.2 | 346.75 | 6.82 | 0.02 |
| 3.06 | + | + | + | + | + | + | NA | + | + | + | + | NA | NA | NA | NA | 42 | -129.2 | 349.07 | 9.14 | 0.01 |
| 3.06 | + | + | + | + | + | + | + | + | NA | NA | + | NA | NA | NA | NA | 44 | -128.6 | 352.67 | 12.74 | 0.00 |
| 3.06 | + | + | + | + | + | + | NA | + | + | + | + | NA | NA | + | NA | 44 | -129 | 353.51 | 13.58 | 0.00 |
| 3.06 | + | + | + | + | + | + | + | + | NA | + | + | NA | NA | NA | NA | 45 | -128.6 | 355.01 | 15.09 | 0.00 |
| 3.06 | + | + | + | + | + | + | + | + | + | NA | + | NA | NA | NA | NA | 46 | -128.6 | 357.29 | 17.36 | 0.00 |
| 3.06 | + | + | + | + | + | + | + | + | + | + | + | NA | NA | NA | NA | 47 | -128.6 | 359.65 | 19.72 | 0.00 |

| **B** | **Value** | **CI (lower)** | **CI (upper)** | **Std.Error** | **DF** | **t-value** | **p-value** | |
|---|---|---|---|---|---|---|---|---|
| Region:Kiel (at 15 C, least dilute) | 3.06 | 2.90 | 3.21 | 0.08 | 13 | 38.24 | <0.001 | *** |

| | | | | | | | | |
|---|---|---|---|---|---|---|---|---|
| Region: Bornholm (at 15 C, least dilute) | 0.54 | 0.32 | 0.76 | 0.11 | 13 | 4.81 | <0.001 | *** |
| Temp18 | 0.42 | 0.21 | 0.64 | 0.11 | 13 | 3.89 | <0.001 | *** |
| Temp22 | 1.05 | 0.82 | 1.28 | 0.12 | 13 | 8.96 | <0.001 | *** |
| Dilution1e-05 | 0.09 | -0.13 | 0.31 | 0.11 | 13 | 0.77 | 0.441 | |
| Dilution1e-04 | 0.10 | -0.11 | 0.30 | 0.11 | 13 | 0.89 | 0.374 | |
| Dilution0.001 | 0.19 | -0.02 | 0.39 | 0.11 | 13 | 1.76 | 0.079 | . |
| Dilution0.01 | -0.01 | -0.21 | 0.20 | 0.11 | 13 | -0.07 | 0.942 | |
| Dilution0.1 | -0.09 | -0.30 | 0.12 | 0.11 | 13 | -0.82 | 0.41 | |
| Region: Bornholm: Temp18 | -0.70 | -1.00 | -0.39 | 0.16 | 13 | -4.51 | <0.001 | *** |
| Region: Bornholm: Temp22 | -1.81 | -2.14 | -1.48 | 0.17 | 13 | -10.78 | <0.001 | *** |
| Region: Bornholm: Dilution1e-05 | -0.32 | -0.64 | 0.00 | 0.16 | 13 | -1.95 | 0.052 | . |
| Region: Bornholm: Dilution1e-04 | -0.22 | -0.53 | 0.09 | 0.16 | 13 | -1.40 | 0.161 | |
| Region: Bornholm: Dilution0.001 | -0.34 | -0.66 | -0.03 | 0.16 | 13 | -2.16 | 0.031 | * |
| Region: Bornholm: Dilution0.01 | -0.37 | -0.68 | -0.06 | 0.16 | 13 | -2.37 | 0.018 | * |
| Region: Bornholm: Dilution0.1 | -0.22 | -0.53 | 0.09 | 0.16 | 13 | -1.38 | 0.169 | |
| Temp18: Dilution1e-05 | -0.18 | -0.49 | 0.13 | 0.16 | 13 | -1.16 | 0.248 | |
| Temp22: Dilution1e-05 | -0.17 | -0.50 | 0.15 | 0.16 | 13 | -1.05 | 0.296 | |
| Temp18: Dilution1e-04 | -0.07 | -0.37 | 0.22 | 0.15 | 13 | -0.49 | 0.623 | |
| Temp22: Dilution1e-04 | -0.41 | -0.73 | -0.09 | 0.16 | 13 | -2.55 | 0.011 | * |
| Temp18: Dilution0.001 | 0.05 | -0.25 | 0.35 | 0.15 | 13 | 0.31 | 0.759 | |
| Temp22: Dilution0.001 | -0.39 | -0.70 | -0.08 | 0.16 | 13 | -2.43 | 0.015 | * |
| Temp18: Dilution0.01 | 0.15 | -0.15 | 0.44 | 0.15 | 13 | 0.98 | 0.327 | |
| Temp22: Dilution0.01 | -0.18 | -0.50 | 0.13 | 0.16 | 13 | -1.15 | 0.25 | |
| Temp18: Dilution0.1 | 0.05 | -0.25 | 0.35 | 0.15 | 13 | 0.34 | 0.731 | |
| Temp22: Dilution0.1 | -0.06 | -0.38 | 0.25 | 0.16 | 13 | -0.39 | 0.694 | |
| Region: Bornholm: Temp18: Dilution1e-05 | 0.70 | 0.26 | 1.13 | 0.22 | 13 | 3.14 | 0.002 | ** |
| Region: Bornholm: Temp22: Dilution1e-05 | 0.90 | 0.44 | 1.37 | 0.24 | 13 | 3.82 | <0.001 | *** |
| Region: Bornholm: Temp18: Dilution1e-04 | 0.46 | 0.03 | 0.89 | 0.22 | 13 | 2.12 | 0.034 | * |

| | | | | | | | | |
|---|---|---|---|---|---|---|---|---|
| Region: Bornholm: Temp22: Dilution1e-04 | 1.43 | 0.98 | 1.88 | 0.23 | 13 | 6.22 | <0.001 | *** |
| Region: Bornholm: Temp18: Dilution0.001 | 0.40 | -0.04 | 0.83 | 0.22 | 13 | 1.80 | 0.072 | . |
| Region: Bornholm: Temp22: Dilution0.001 | 1.73 | 1.28 | 2.19 | 0.23 | 13 | 7.54 | <0.001 | *** |
| Region: Bornholm: Temp18: Dilution0.01 | 0.69 | 0.27 | 1.11 | 0.22 | 13 | 3.20 | 0.001 | ** |
| Region: Bornholm: Temp22: Dilution0.01 | 1.49 | 1.04 | 1.94 | 0.23 | 13 | 6.53 | <0.001 | *** |
| Region: Bornholm: Temp18: Dilution0.1 | 0.63 | 0.20 | 1.06 | 0.22 | 13 | 2.87 | 0.004 | ** |
| Region: Bornholm: Temp22: Dilution0.1 | 1.43 | 0.98 | 1.88 | 0.23 | 13 | 6.23 | <0.001 | *** |

**Table S4: Model selection (A) output (B) for investigating the effect of dilution (abbreviated to D), assay temperature (abbreviated T), region (abbreviated R), and season (abbreviated S) on cell diameter (µm) at carrying capacity.**
In the mixed model, D (from 1 – highest richness to 1e-05 – lowest richness), T (15ºC,18ºC, 22ºC), R (Kiel Area, Bornholm Basin), and S (spring, summer) selection regimes, i.e. nutrient (low nutrient and replete), were fitted as fixed effects. Stations were treated as a random factor. Technical replicates were not fitted. The best model (highlighted in bold) is the model with the smallest AICc, where delta AICc to the next best model is >2. df for degrees of freedom; logLik for log likelihood ratio. : indicates an interaction term. We display only the first 10 models for clarity.
The global model formula was lme.formula(sizeum~D*R*S*T, random=~1|bio.stat.id, data=dataframe.sizeum, method="ML) . The model used for the model output table was refitted with REML and read lme.formula(sizeum ~D*R*T, random=~1|bio.stat.id, data=dataframe. sizeum, method="REML). In the model output table, CI are the 95% confidence intervals, DF are degrees of freedom. Values other than the first value (Kiel sample at 15ºC with the lowest dilution, i.e. highest diversity) need to be added to the first value to obtain the predicted trait value.

| A) Intercept | D | T | R | S | D:T | D:R | D:S | T:R | S:T | R:S | D:T:R | D:T:S | D:R:S | T:R:S | D:T:R:S | df | logLik | AICc | Δ | weight |
|---|---|---|---|---|---|---|---|---|---|---|---|---|---|---|---|---|---|---|---|---|
| **2.92** | **+** | **+** | **+** | **NA** | **+** | **+** | **NA** | **+** | **NA** | **NA** | **+** | **NA** | **NA** | **NA** | **NA** | **38** | **-863.16** | **1807.83** | **0** | **0.65** |
| 2.9 | + | + | + | + | + | + | NA | + | NA | NA | + | NA | NA | NA | NA | 39 | -863.06 | 1809.92 | 2.1 | 0.23 |
| 2.9 | + | + | + | + | + | + | NA | + | NA | + | + | NA | NA | NA | NA | 40 | -863.03 | 1812.18 | 4.35 | 0.07 |
| 2.96 | + | + | + | + | + | + | NA | + | + | NA | + | NA | NA | NA | NA | 41 | -862.61 | 1813.66 | 5.83 | 0.03 |
| 2.95 | + | + | + | + | + | + | NA | + | + | + | + | NA | NA | NA | NA | 42 | -862.57 | 1815.91 | 8.08 | 0.01 |
| 2.93 | + | + | + | + | + | + | NA | + | + | + | + | NA | NA | + | NA | 44 | -861.26 | 1817.97 | 10.14 | 0 |
| 2.89 | + | + | + | + | + | + | + | + | NA | NA | + | NA | NA | NA | NA | 44 | -862.29 | 1820.02 | 12.19 | 0 |
| 2.88 | + | + | + | + | + | + | + | + | NA | + | + | NA | NA | NA | NA | 45 | -862.26 | 1822.32 | 14.49 | 0 |
| 2.94 | + | + | + | + | + | + | + | + | + | NA | + | NA | NA | NA | NA | 46 | -861.83 | 1823.82 | 15.99 | 0 |
| 2.93 | + | + | + | + | + | + | + | + | + | + | + | NA | NA | NA | NA | 47 | -861.79 | 1826.12 | 18.29 | 0 |

| B) | Value | CI (lower) | CI (upper) | Std.Error | DF | t-value | p-value |
|---|---|---|---|---|---|---|---|

| | | | | | | | | |
|---|---|---|---|---|---|---|---|---|
| Region:Kiel (at 15 C, least dilute) | 2.92 | 3.39 | 4.46 | 0.27 | 13 | 14.42 | <0.001 | *** |
| Region: Bornholm (at 15 C, least dilute) | 1.14 | 1.35 | 2.93 | 0.40 | 13 | 5.33 | <0.001 | *** |
| Temp18 | 0.73 | -0.04 | 1.49 | 0.39 | 13 | 1.87 | 0.062 | . |
| Temp22 | 1.17 | 1.34 | 3.00 | 0.42 | 13 | 5.16 | <0.001 | *** |
| Dilution1e-05 | 0.75 | -0.04 | 1.54 | 0.40 | 13 | 1.86 | 0.064 | . |
| Dilution1e-04 | 0.40 | -0.35 | 1.15 | 0.38 | 13 | 1.06 | 0.292 | |
| Dilution0.001 | 0.17 | -0.57 | 0.92 | 0.38 | 13 | 0.46 | 0.643 | |
| Dilution0.01 | 0.15 | -0.59 | 0.89 | 0.38 | 13 | 0.40 | 0.687 | |
| Dilution0.1 | 0.42 | -0.34 | 1.19 | 0.39 | 13 | 1.09 | 0.275 | |
| Region: Bornholm: Temp18 | -0.01 | -1.10 | 1.08 | 0.56 | 13 | -0.02 | 0.988 | |
| Region: Bornholm: Temp22 | -2.53 | -3.71 | -1.35 | 0.60 | 13 | -4.21 | <0.001 | *** |
| Region: Bornholm: Dilution1e-05 | -0.75 | -1.89 | 0.40 | 0.58 | 13 | -1.28 | 0.2 | |
| Region: Bornholm: Dilution1e-04 | -0.25 | -1.35 | 0.86 | 0.56 | 13 | -0.44 | 0.662 | |
| Region: Bornholm: Dilution0.001 | -0.36 | -1.48 | 0.76 | 0.57 | 13 | -0.63 | 0.527 | |
| Region: Bornholm: Dilution0.01 | -2.73 | -3.83 | -1.63 | 0.56 | 13 | -4.87 | <0.001 | *** |
| Region: Bornholm: Dilution0.1 | -3.30 | -4.41 | -2.18 | 0.57 | 13 | -5.81 | <0.001 | *** |
| Temp18: Dilution1e-05 | -0.73 | -1.83 | 0.37 | 0.56 | 13 | -1.31 | 0.191 | |
| Temp22: Dilution1e-05 | -1.69 | -2.85 | -0.54 | 0.59 | 13 | -2.87 | 0.004 | ** |
| Temp18: Dilution1e-04 | -0.63 | -1.69 | 0.44 | 0.54 | 13 | -1.16 | 0.247 | |
| Temp22: Dilution1e-04 | -1.46 | -2.60 | -0.33 | 0.58 | 13 | -2.54 | 0.011 | * |
| Temp18: Dilution0.001 | -0.57 | -1.64 | 0.50 | 0.55 | 13 | -1.04 | 0.299 | |
| Temp22: Dilution0.001 | -1.36 | -2.49 | -0.24 | 0.57 | 13 | -2.38 | 0.018 | * |
| Temp18: Dilution0.01 | -0.48 | -1.54 | 0.57 | 0.54 | 13 | -0.90 | 0.368 | |
| Temp22: Dilution0.01 | -1.40 | -2.53 | -0.27 | 0.57 | 13 | -2.44 | 0.015 | * |
| Temp18: Dilution0.1 | -0.47 | -1.54 | 0.60 | 0.55 | 13 | -0.87 | 0.387 | |
| Temp22: Dilution0.1 | -2.10 | -3.24 | -0.96 | 0.58 | 13 | -3.62 | <0.001 | *** |
| Region: Bornholm: Temp18: Dilution1e-05 | 0.73 | -0.82 | 2.29 | 0.79 | 13 | 0.92 | 0.356 | |
| Region: Bornholm: Temp22: Dilution1e-05 | 1.69 | 0.03 | 3.36 | 0.85 | 13 | 2.00 | 0.046 | * |

| | | | | | | | | |
|---|---|---|---|---|---|---|---|---|
| Region: Bornholm: Temp18: Dilution1e-04 | 0.10 | -1.43 | 1.62 | 0.78 | 13 | 0.12 | 0.903 | |
| Region: Bornholm: Temp22: Dilution1e-04 | 1.43 | -0.18 | 3.04 | 0.82 | 13 | 1.74 | 0.083 | . |
| Region: Bornholm: Temp18: Dilution0.001 | -0.37 | -1.92 | 1.19 | 0.79 | 13 | -0.46 | 0.643 | |
| Region: Bornholm: Temp22: Dilution0.001 | 1.29 | 0.67 | 3.90 | 0.82 | 13 | 2.78 | 0.006 | ** |
| Region: Bornholm: Temp18: Dilution0.01 | 1.66 | 0.14 | 3.18 | 0.77 | 13 | 2.14 | 0.032 | * |
| Region: Bornholm: Temp22: Dilution0.01 | 3.05 | 2.45 | 5.65 | 0.82 | 13 | 4.97 | <0.001 | *** |
| Region: Bornholm: Temp18: Dilution0.1 | 1.85 | 0.30 | 3.40 | 0.79 | 13 | 2.35 | 0.019 | * |
| Region: Bornholm: Temp22: Dilution0.1 | 3.45 | 3.84 | 7.06 | 0.82 | 13 | 6.65 | <0.001 | *** |

**Table S5: Model selection (A) output (B) for investigating the effect of dilution (abbreviated to D), assay temperature (abbreviated T), region (abbreviated R), and season (abbreviated S) on Net Photosynthesis rates (µmol $O_2$ per cell and hour - displayed as LOG10 values for brevity) during exponential growth.**
In the mixed model, D (from 1 – highest richness to 100000– lowest richness), T (15ºC,18ºC, 22ºC), R (Kiel Area, Bornholm Basin), and S (spring, summer) selection regimes, i.e. nutrient (low nutrient and replete), were fitted as fixed effects. Stations were treated as a random factor. Technical replicates were not fitted. The best model (highlighted in bold) is the model with the smallest AICc, where delta AICc to the next best model is >2. df for degrees of freedom; logLik for log likelihood ratio. : indicates an interaction term. We display only the first 10 models for clarity.
The global model formula was lme.formula(sizeum~D*R*S*T, random=~1|bio.stat.id, data=dataframe.sizeum, method="ML) . The model used for the model output table was refitted with REML and read lme.formula(sizeum ~R*T +D, random=~1|bio.stat.id, data=dataframe. sizeum, method="REML). In the model output table, CI are the 95% confidence intervals, DF are degrees of freedom. Values other than the first value (Kiel sample at 15ºC with the lowest dilution, i.e. highest diversity) need to be added to the first value to obtain the predicted trait value.

| Intercept | T | D | R | S | T*D | T*R | T*S | D*R | D*S | R*S | T*D*R | T*D*S | T*R*S | D*R*S | T*D*R*S | df | logLik | AICc | delta | weight |
|---|---|---|---|---|---|---|---|---|---|---|---|---|---|---|---|---|---|---|---|---|
| **-7.02E-09** | **+** | **+** | **+** | **NA** | **NA** | **+** | **NA** | **+** | **NA** | **NA** | **NA** | **NA** | **NA** | **NA** | **NA** | **18** | **3397.38** | **-6755.28** | **0.00** | **0.36** |
| -7.01E-09 | + | + | + | + | NA | + | + | + | NA | NA | NA | NA | NA | NA | NA | 21 | 3400.32 | -6757.35 | 2.08 | 0.18 |
| -6.93E-09 | + | + | + | + | NA | + | NA | + | NA | NA | NA | NA | NA | NA | NA | 19 | 3397.38 | -6752.87 | 2.40 | 0.11 |
| 8.07E-09 | + | NA | + | NA | NA | + | NA | NA | NA | NA | NA | NA | NA | NA | NA | 8 | 3384.71 | -6752.72 | 2.56 | 0.10 |
| -6.19E-09 | + | + | + | + | NA | + | + | + | NA | + | NA | NA | NA | NA | NA | 22 | 3400.37 | -6751.51 | 3.77 | 0.05 |
| 8.08E-09 | + | NA | + | + | NA | + | + | NA | NA | NA | NA | NA | NA | NA | NA | 11 | 3387.32 | -6751.35 | 3.93 | 0.05 |
| -6.12E-09 | + | + | + | + | NA | + | NA | + | NA | + | NA | NA | NA | NA | NA | 20 | 3397.43 | -6750.56 | 4.72 | 0.03 |
| 8.16E-09 | + | NA | + | + | NA | + | NA | NA | NA | NA | NA | NA | NA | NA | NA | 9 | 3384.71 | -6750.55 | 4.73 | 0.03 |
| -6.99E-09 | + | + | + | + | NA | + | + | + | NA | + | NA | NA | + | NA | NA | 24 | 3402.21 | -6750.14 | 5.14 | 0.03 |
| 8.90E-09 | + | NA | + | + | NA | + | + | NA | NA | + | NA | NA | NA | NA | NA | 12 | 3387.37 | -6749.21 | 6.07 | 0.02 |
| 8.97E-09 | + | NA | + | + | NA | + | NA | NA | NA | + | NA | NA | NA | NA | NA | 10 | 3384.76 | -6748.45 | 6.83 | 0.01 |

| **B)** | **Value** | **CI (lower)** | **CI (upper)** | **Std.Error** | **DF** | **t-value** | **p-value** |
|---|---|---|---|---|---|---|---|
| Region:Kiel (at 15 C, least dilute) | 1.00E-08 | -7.07E-09 | 2.71E-08 | 8.66E-09 | 200 | 1.16 | 0.249 |

| | | | | | | | | |
|---|---|---|---|---|---|---|---|---|
| Region: Bornholm (at 15 C, least dilute) | -6.28E-09 | -2.41E-08 | 1.16E-08 | 9.05E-09 | 200 | -0.69 | 0.489 | |
| Temp18 | 2.43E-08 | 6.44E-09 | 4.21E-08 | 9.05E-09 | 200 | 2.68 | 0.008 | ** |
| Temp22 | 9.39E-08 | 7.60E-08 | 1.12E-07 | 9.05E-09 | 200 | 10.37 | <0.001 | *** |
| Dilution10 | -5.12E-09 | -2.30E-08 | 1.27E-08 | 9.05E-09 | 200 | -0.57 | 0.572 | |
| Dilution100 | -7.96E-09 | -2.58E-08 | 9.88E-09 | 9.05E-09 | 200 | -0.88 | 0.38 | |
| Dilution1000 | -6.64E-09 | -2.45E-08 | 1.12E-08 | 9.05E-09 | 200 | -0.73 | 0.464 | |
| Dilution10000 | 9.95E-09 | -7.90E-09 | 2.78E-08 | 9.05E-09 | 200 | 1.10 | 0.273 | |
| Dilution100000 | -1.89E-09 | -1.97E-08 | 1.60E-08 | 9.05E-09 | 200 | -0.21 | 0.835 | |
| Region: Bornholm: Temp18 | -1.31E-08 | -3.83E-08 | 1.22E-08 | 1.28E-08 | 200 | -1.02 | 0.308 | |
| Region: Bornholm:Temp22 | -7.33E-08 | -9.86E-08 | -4.81E-08 | 1.28E-08 | 200 | -5.73 | <0.001 | *** |
